# RapMap: A Rapid, Sensitive and Accurate Tool for Mapping RNA-seq Reads to Transcriptomes

**DOI:** 10.1101/029652

**Authors:** Avi Srivastava, Hirak Sarkar, Nitish Gupta, Rob Patro

## Abstract

**Motivation:** The alignment of sequencing reads to a transcriptome is a common and important step in many RNA-seq analysis tasks. When aligning RNA-seq reads directly to a transcriptome (as is common in the *de novo* setting or when a trusted reference annotation is available), care must be taken to report the potentially large number of multi-mapping locations per read. This can pose a substantial computational burden for existing aligners, and can considerably slow downstream analysis.

**Results:** We introduce a novel concept, quasi-mapping, and an efficient algorithm implementing this approach for mapping sequencing reads to a transcriptome. By attempting only to report the potential loci of origin of a sequencing read, and not the base-to-base alignment by which it derives from the reference, `RapMap`— our tool implementing quasi-mapping— is capable of *mapping* sequencing reads to a target transcriptome substantially faster than existing alignment tools. The algorithm we employ to implement quasi-mapping uses several efficient data structures and takes advantage of the special structure of shared sequence prevalent in transcriptomes to rapidly provide highly-accurate mapping information. We demonstrate how quasi-mapping can be successfully applied to the problems of transcript-level quantification from RNA-seq reads and the clustering of contigs from *de novo* assembled transcriptomes into biologically-meaningful groups.

**Availability:** `RapMap` is implemented in `C++11` and is available as open-source software, under GPL v3, at https://github.com/COMBINE-lab/RapMap.

## 1 Introduction

The bioinformatics community has put tremendous effort into building a wide array of different tools to solve the read-alignment problem efficiently. These tools use many different strategies to quickly find potential alignment locations for reads; for example, Bowtie (Langmead *et al.*, 2009), `Bowtie 2` (Langmead and Salzberg, 2012), BWA (Li and Durbin, 2009) and BWA-mem (Li, 2013) use variants of the FM-index, while tools like the Subread aligner (Liao *et al.*, 2013), Maq (Li *et al.*, 2008) and MrsFast (Hach *et al.*, 2010) use k-mer-based indices to help align reads efficiently. Because read alignment is such a ubiquitous task, the goal of such tools is often to provide accurate results as quickly as possible. Indeed, recent alignment tools like `STAR` (Dobin *et al.*, 2013) demonstrate that rapid alignment of sequenced reads is possible, and tools like HISAT (Kim *et al.*, 2015) demonstrate that this speed can be achieved with only moderate memory usage. When reads are aligned to a collection of reference sequences that share a substantial amount of sub-sequence (near or exact repeats), a single read can have many potential alignments, and considering all such alignment can be crucial for downstream analysis (e.g. considering all alignment locations for a read within a transcriptome for the purpose of quantification Li and Dewey (2011), or when attempting to cluster *de novo* assembled contigs by shared multi-mapping reads (Davidson and Oshlack, 2014)). However, reporting multiple potential alignments for each read is a difficult task, and tends to substantially slow down even very efficient alignment tools.

Yet, in many cases, all of the information provided by the alignments is not necessary. For example, in the transcript analysis tasks mentioned above, simply the knowledge of the transcripts and positions to which a given read maps well is sufficient to answer the questions being posed. In support of such “analysis-oriented” computation, we propose a novel concept, called quasi-mapping, and an efficient algorithm implementing quasi-mapping (exposed in the software tool `RapMap`) to solve the problem of mapping sequenced reads to a target transcriptome. This algorithm is *considerably* faster than state-of-the-art aligners, and achieves its impressive speed by exploiting the structure of the transcriptome (without requiring an annotation), and eliding the computation of full-alignments (e.g. `CIGAR` strings). Further, our algorithm produces mappings that meet or exceed the accuracy of existing popular aligners under different metrics of accuracy. Finally, we demonstrate how the mappings produced by `RapMap` can be used in the downstream analysis task of transcript-level quantification from RNA-seq data, by modifying the Sailfish (Patro *et al.*, 2014) tool to take advantage of quasi-mappings, as opposed to raw k-mer counts, for transcript quantification. We also demonstrate how quasi-mappings can be used to effectively cluster contigs from *de novo* assemblies. We show that the resulting clusterings are of comparable or superior accuracy to those produced by recent methods such as CORSET (Davidson and Oshlack, 2014), but that they can be computed *much* more quickly using quasi-mapping.

## 2 Methods

The quasi-mapping concept, implemented in the tool `RapMap`, is a new mapping technique to allow the rapid and accurate mapping of sequenced fragments (single or paired-end reads) to a target transcriptome. `RapMap` exploits a combination of data structures — a hash table, suffix array (SA), and efficient rank data structure. It takes into account the special structure present in transcriptomic references, as exposed by the suffix array, to enable ultra-fast and accurate determination of the likely loci of origin of a sequencing read. Rather than a standard alignment, quasi-mapping produces what we refer to as fragment *mapping* information. In particular, it provides, for each query (fragment), the reference sequences (transcripts), strand and position from which the query may have likely originated. In many cases, this mapping information is sufficient for downstream analysis. For example, tasks like transcript quantification, clustering of *de novo* assembled transcripts, and filtering of potential target transcripts can be accomplished with this mapping information. However, this method does not compute the base-to-base alignment between the query and reference. Thus, such mappings may not be appropriate in every situation in which alignments are currently used (e.g. variant detection).

We note here that the concept of quasi-mapping shares certain motivations with the notions of lightweight-alignment (Patro *et al.*, 2015) and pseudo-alignment (Bray *et al.*, 2015). Yet, all three concepts — and the algorithms and data structures used to implement them — are distinct and, in places, substantially different. Lightweight-alignment scores potential matches based on approximately consistent chains of super-maximal exact matches shared between the query and targets. Therefore, it typically requires some more computation than the other methods, but allows the reporting of a score with each returned mapping and a more flexible notion of matching. Pseudo-alignment, as implemented in `Kallisto`, refers only to the process of finding *compatible* targets for reads by determining approximately matching paths in a colored De Bruijn graph of a pre-specified order. Among compatible targets, extra information concerning the mapping (e.g. position and orientation) can be extracted *post-hoc*, but this requires extra processing, and the resulting mapping is no longer technically a pseudo-alignment. Quasi-mapping seeks to find the *best* mappings (targets and positions) for each read, and does so (approximately) by finding minimal collections of dynamically-sized, right-maximal matching contexts between target and query positions. The algorithm for quasi-mapping that we describe below achieves this using a combination of a k-mer lookup table and a generalized suffix array. While each of these approaches provide some insight into the problems of alignment and mapping, they represent distinct concepts and exhibit unique characteristics in terms of speed and accuracy, as demonstrated below^1^.

*An algorithm for Quasi-mapping* The algorithm we use for quasi-mapping makes use of two main data structures, the generalized suffix array (Manber and Myers, 1993) `SA`[*T*] of the transcriptome *T*, and a hash table *h* mapping each k-mer occurring in *T* to its suffix array interval (by default *k* = 31). Additionally, we must maintain the original text *T* upon which the suffix array was constructed, and the name and length of each of the original transcript sequences. *T* consists of a string in which all transcript sequences are joined together with a special separator character. Rather than designating a separate terminator $_i_ for each reference sequence in the transcriptome, we make use of a single separator $, and maintain an auxiliary rank data structure which allows us to map from an arbitrary position in the concatenated text to the index of the reference transcript in which it appears. We use the rank9b algorithm and data structure of Vigna (2008) to perform the rank operation quickly.

Quasi-mapping determines the mapping locations for a query read *r* through repeated application of (1) determining the next hash table k-mer that starts past the current query position, (2) computing the maximum mappable prefix (MMP) of the query beginning with this k-mer, and then (3) determining the next informative position (NIP) by performing a longest common prefix (LCP) query on two specifically chosen suffixes in the suffix array.

The algorithm begins by hashing the k-mers of *r*, from left-to-right (a symmetric procedure can be used for mapping the reverse-complement of a read), until some k-mer *k_i_* — the k-mer starting at position *i* within the read — is present in *h* and maps to a valid suffix array interval. We denote this interval as 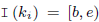. Because of the lexicographic order of the suffixes in the suffix array, we immediately know that this k-mer is a prefix of all of the suffixes appearing in the given interval. However, it may be possible to extend this match to some longer substring of the read beginning with *k_i_*. In fact, the longest substring of the read that appears in the reference and is prefixed by *k_i_* is exactly the maximum mappable prefix (MMP) (Dobin *et al.*, 2013) of the suffix of the read beginning with *k_i_*. We call this maximum mappable prefix MMP_i_, and note that it can be found using a slight variant of the standard suffix array binary search (Manber and Myers, 1993) algorithm. For speed and simplicity, we implement the “simple accelerant” binary search variant of Gusfield (1997). Since we know that any substring that begins with *k_i_* must reside in the interval [*b, e*), we can restrict the MMP*_i_* search to this region of the suffix array, which is typically very small.

After determining the length of MMP*_i_* within the read, one could begin the search for the next mappable suffix array interval at the position following this MMP. However, though the current substring of the read will differ from all of the reference sequence suffixes at the base following MMP*_i_*, the suffixes occurring at the lower and upper bounds of the suffix array interval corresponding to MMP*_i_* may not differ from each other (see Figure 1). That is, if 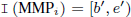 is the suffix array interval corresponding to MMP*_i_*, it is possible that 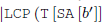, 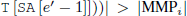. In this case, it is most likely that the read and the reference sequence bases following MMP*_i_* disagree as the result of a sequencing error, not because the (long) MMP discovered between the read and reference is a spurious match. Thus, beginning the search for the next MMP at the subsequent base in the read may not be productive, since the matches for this substring of the query may not be informative — that is, such a search will likely return the same (relative) positions and set of transcripts. To avoid querying for such substrings, we define and make use of the notion of the next informative position (NIP). For a maximum mappable prefix MMP*_i_*, with 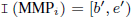, we define NIP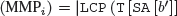, 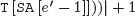. Intuitively, the next informative position of prefix MMP*_i_* is designed to return the next position in the query string where a suffix array search is likely to yield a set of transcripts different from those contained in I(MMP*_i_*). To compute the longest common prefix between two suffixes when searching for the NIP, we use the “direct min” algorithm of Ilie *et al*. (2010). We found this to be the fastest approach. Additionally, it doesn’t require the maintenance of an LCP array or other auxiliary tables aside from the standard suffix array.

**Fig 1:**
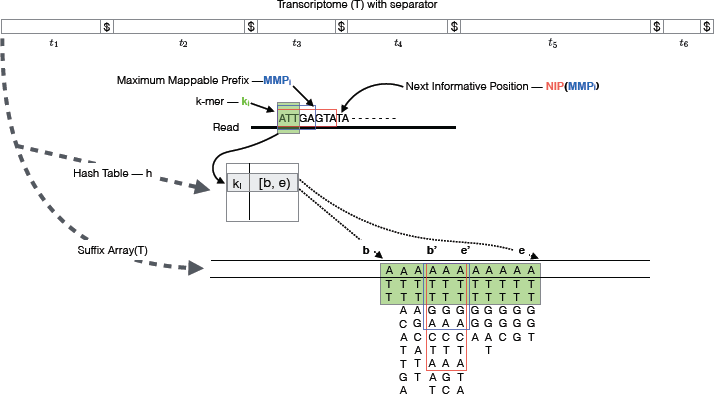
The transcriptome (consisting of transcripts *t*_1_,…, *t*_6_) is converted into a $-separated string, *T*, upon which a suffix array, `SA`[*T*], and a hash table, *h*, are constructed. The mapping operation begins with a k-mer (here, *k* = 3) mapping to an interval [*b, e*) in `SA`[*T*]. Given this interval and the read, MMP*_i_* and `NIP`(MMP*_i_*) are calculated as described in section 2. The search for the next hashable k-mer begins *k* bases before `NIP`(MMP*_i_*).

Given the definitions we have explained above, we can summarize the quasi-mapping procedure as follows (an illustration of the mapping procedure is provided in Figure 1). First, a read is scanned from left to right (a symmetric procedure can be used for mapping the reverse-complement of a read) until a k-mer *k_i_* is encountered that appears in *h*. A lookup in h returns the suffix array interval I (*k_i_*) corresponding to the substring of the read consisting of this k-mer. Then, the procedure described above is used to compute MMP*_i_* and 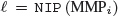. The search procedure then advances to position *i* + *l* — *k* in the read, and again begins hashing the k-mers it encounters. This process of determining the MMP and NIP of each processed k-mer and advancing to the next informative position in the read continues until the next informative position exceeds position *l_r_* — *k* where *l_r_* is the length of the read *r*. The result of applying this procedure to a read is a set 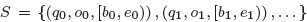 of query positions, MMP orientations, and suffix array intervals, with one such triplet corresponding to each MMP.

The final set of mappings is determined by a consensus mechanism. Specifically, the algorithm reports the set of transcripts that appear in every suffix array interval appearing in *S*. These transcripts, and the corresponding strand and location on each, are reported as *quasi-mappings* of this read. These mappings are reported in a `samtools`-compatible format in which the relevant information (e.g. target id, position, strand, pair status) is computed from the mapping. In the next section, we analyze how this algorithm for quasi-mapping, as described above compares to other aligners in terms of speed and mapping accuracy.

## 3 Mapping speed and accuracy

To test the practical performance of quasi-mapping, we compared `RapMap` against a number of existing tools, and analyzed both the speed and accuracy of these tools on synthetic and experimental data. Benchmarking was performed against the popular aligners `Bowtie 2` (Langmead and Salzberg, 2012) (v2.2.6) and `STAR` (Dobin *et al.*, 2013) (v2.5.0c) and the recently-introduced pseudo-alignment procedure used in the quantification tool `Kallisto` (Bray *et al.*, 2015) (v0.42.4). All experiments were scripted using Snakemake (Köster and Rahmann, 2012) and performed on a 64-bit linux server with 256GB of RAM and 4 x 6-core Intel Xeon E5-4607 v2 CPUs running at 2.60GHz. Wall-clock time was recorded using the `time` command.

In our testing we find that `Bowtie 2` generally performs well in terms of reporting the true read origin among its set of multi-mapping locations. However, it takes considerably longer and tends to return a larger set of multi-mapping locations than the other methods. In comparison to `Bowtie 2, STAR` is *substantially* faster but somewhat less accurate. `RapMap` achieves accuracy comparable or superior to `Bowtie 2`, while simultaneously being much faster than even `STAR`. `Kallisto` is similar to (slightly slower than) `RapMap` in terms of single-threaded speed, and exhibits accuracy very similar to that of `STAR`. For both `RapMap` and `Kallisto`, simply writing the output to disk tends to dominate the time required for large input files with significant multi-mapping. This is due, in part, to the verbosity of the standard `SAM` format in which results are reported, and suggests that it may be worth developing a more efficient and succinct output format for mapping information.

### 3.1 Speed and accuracy on synthetic data

To test the accuracy of different mapping and alignment tools in a scenario where we know the true origin of each read, we generated data using the Flux Simulator (Griebel *et al*, 2012). This synthetic dataset was generated for the human transcriptome from an annotation taken from the ENSEMBL (Cunningham *et al*, 2015) database consisting of 86,090 transcripts corresponding to protein-coding genes. The dataset consists of ~ 48 million 76 base pair, paired-end reads. The detailed parameters used for the Flux Simulator can be found in Appendix 1.2.

When benchmarking these methods, reads were aligned directly to the transcriptome, rather than to the genome. This was done because we wish to benchmark the tools in a manner that is applicable when the reference genome may not even be known (e.g. in *de novo* transcriptomics). The parameters of `STAR` (see Appendix 1.1) were adjusted appropriately for this purpose (e.g. to dis-allow introns etc.). Similarly, `Bowtie 2` was also used to align reads directly to the target transcriptome; the parameters for `Bowtie 2` are given in Appendix 1.1.

#### 3.1.1 Mapping speed

We wish to measure, as directly as possible, just the time required by the mapping algorithms of the different tools. Thus, when benchmarking the runtime of different methods, we do not save the resulting alignments to disk. Further, to mitigate the effect of “outliers” (a small number of reads which map to a very large number of low-complexity reference positions), we bound the number of different transcripts to which a read can map to be 200. Additionally, we have also benchmarked `Kallisto`, but have not included the results in Figure 2, as the software, unlike the other methods, does not allow multi-threaded execution if mappings are being reported. Thus, we ran `Kallisto` with a single thread, using the -`pseudobam` flag and redirecting output to `/dev/null` to avoid disk overhead. `Kallisto` requires 17.87m tomapthe48M simulated reads, which included <1m of quantification time. By comparison, RapMap required 11.65m to complete with a single thread.

**Fig 2:**
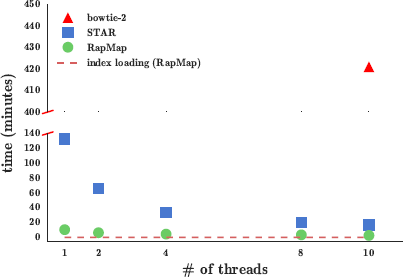
The time taken by `Bowtie 2`, `STAR` and `RapMap` to process the synthetic data using varying numbers of threads. `RapMap` processes the data substantially faster than the other tools, while providing results of comparable or better accuracy.

Finally, we note `Kallisto`, `STAR` and `RapMap` require 2-3× the memory of `Bowtie 2`, but all of the methods tested here exhibit reasonable memory usage. The synthetic set of 48 million reads can be mapped to an index of the entire human transcriptome on a typical laptop with 8 GB of RAM.

As Figure 2 illustrates, `RapMap` out-performs both `Bowtie 2` and `STAR` in terms of speed by a substantial margin, and finishes mapping the reads with a single thread faster than `STAR` and `Bowtie 2` with 10 threads. We consider varying the number of threads used by `RapMap` and `STAR` to demonstrate how performance scales with the number of threads provided. On this dataset, `RapMap` quickly approaches peak performance after using only a few threads. We believe that this is not due to limits on the scalability of `RapMap`, but rather because the process is so quick that, for a dataset of this size, simply reading the index constitutes a large (and growing) fraction of the total runtime (dotted line) as the number of threads is increased. Thus, we believe that the difference in runtime between `RapMap` and the other methods may be even larger for datasets consisting of a very large number of reads, where the disk can reach peak efficiency and the multi-threaded input parser (we use the parser from the Jellyfish (Marçais and Kingsford, 2011) library) can provide input to `RapMap` quickly enough to make use of alarger number of threads. Since running `Bowtie 2` with each potential number of threads on this dataset is very time-consuming, we only consider `Bowtie 2`’s runtime using 10 threads.

#### 3.1.2 Mapping accuracy

Since the Flux Simulator records the true origin of each read, we make use of this information as ground truth data to assess the accuracy of different methods. However, since a single read may have multiple, equally-good alignments with respect to the transcriptome, care must be taken in defining accuracy-related terms appropriately. A read is said to be correctly mapped by a method (a true positive) if the set of transcripts reported by the mapper for this read contains the true transcript. A read is said to be incorrectly mapped by a method (a false positive) if it is mapped to some set of 1 or more transcripts, none of which are the true transcript of origin. Finally, a read is considered to be incorrectly un-mapped by a method (a false negative) if the method reports no mappings, but the transcript of origin is in the reference. Given these definitions, we report precision, recall, F1-Score and false discovery rate (FDR) in Table 1 using the standard definitions of these metrics. Additionally, we report the average number of “hits-per-read” (hpr) returned by each of the methods. Ideally, we want a method to return the smallest set of mappings that contains the true read origin. However, under the chosen definition of a true positive mapping, the number of reported mappings is not taken into account, and a result is considered a true positive so long as it contains the actual transcript of origin. The hpr metric allows one to assess how many *extra* mappings, on average, are reported by a particular method.

**Table 1.**
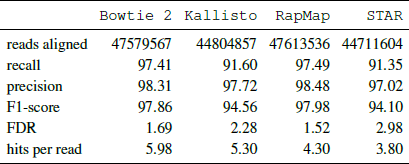
Accuracy of aligners/mappers under different metrics.

|  | Bowtie 2 | Kallisto | RapMap | STAR |
| --- | --- | --- | --- | --- |
| reads aligned | 47579567 | 44804857 | 47613536 | 44711604 |
| recall | 97.41 | 91.60 | 97.49 | 91.35 |
| precision | 98.31 | 97.72 | 98.48 | 97.02 |
| F1-score | 97.86 | 94.56 | 97.98 | 94.10 |
| FDR | 1.69 | 2.28 | 1.52 | 2.98 |
| hits per read | 5.98 | 5.30 | 4.30 | 3.80 |

As expected, `Bowtie 2`— perhaps the most common method of directly mapping reads to transcriptomes — performs very well in terms of precision and recall. However, we find that `RapMap` yields very similar (in fact, slightly better) precision and recall. `STAR` and `Kallisto` obtain similar precision to `Bowtie 2` and `RapMap`, but have lower recall. `STAR` and `Kallisto` perform similarly in general, though `Kallisto` achieves a lower (better) FDR than `STAR`. Taking the F1-score as a summary statistic, we observe that all methods perform reasonably well, and that, in general, alignment-based methods do not seem to be more accurate than mapping-based methods. We also observe that `RapMap` yields very accurate mapping results that match or exceed those of `Bowtie 2`.

Additionally, we tested the impact of noisy reads (i.e. reads not generated from the indexed reference) on the accuracy of the different mappers and aligners. To create these background reads, we use a model inspired by (Gilbert *et al.*, 2004), in which reads are sampled from nascent, un-spiced transcripts. The details of this experiment are included in Appendix 1.3.

### 3.2 Speed and concordance on experimental data

We also explore the concordance of `RapMap` with different mapping and alignment approaches using experimental data from the study of Cho *et al.* (2014) (NCBI GEO accession `SRR1293902`). The sample consists of ∼ 26 million 75 base-pair, paired-end reads sequenced on an Illumina HiSeq.

Since we do not know the true origin of each read, we have instead examined the agreement between the different tools (see Figure 3). Intuitively, two tools agree on the mapping locations of a read if they align / map this read to the same subset of the reference transcriptome (i.e. the same set of transcripts). More formally, we define the elements of our universe, *U*, to be tuples consisting of a read identifier and the set of transcripts returned by a particular tool. For example, if, for read *r_i_*, tool *A* returns alignments to transcripts 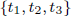, then 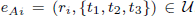. Similarly, if tool *B* maps read *r_i_* to transcripts 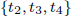 then 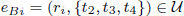. Here, tools *A* and *B* do not agree on the mapping of read *r_i_*. Given a universe *u* thusly-defined, we can employ the normal notions of set intersection and difference to explore how different subsets of methods agree on the mapping locations of the sequenced reads. These concordance results are presented in Figure 3, which uses a bar plot to show the size of each set of potential intersections between the results of the tools we consider. In Figure 3 the dot matrix below the bar plot identifies the tools whose results are intersected to produce the corresponding bar. Tools producing mappings and alignments are denoted with black and red dots and bars respectively. The left bar plot shows the size of the unique tuples produced by each tool (alignments / mappings that do not match with any other tool). The right bar plot shows the total number of tuples produced by each tool, and well as the concordance among all different subsets of tools.

**Fig 3:**
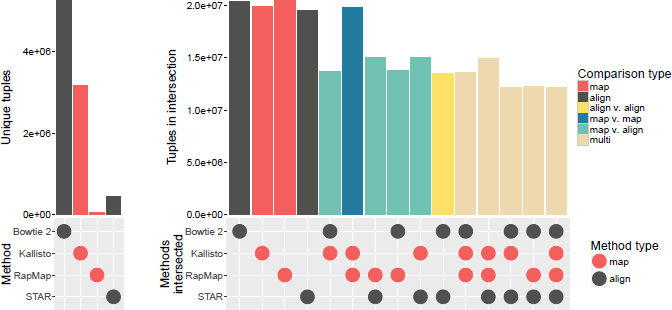
Mapping agreement between subsets of `Bowtie 2`, `STAR`, `Kallisto` and `RapMap`.

Under this measure of agreement, `RapMap` and `Kallisto` appear to agree on the exact same transcript assignments for the largest number of reads. Further, `RapMap` and `Kallisto` have the largest pairwise agreements with the aligners (`STAR` and `Bowtie 2`) — that is, the traditional aligners exactly agree more often with these tools than with each other. It is important to note that one possible reason we see (seemingly) low agreement between `Bowtie 2` and other methods is because the transcript alignment sets reported by `Bowtie 2` are generally larger (i.e. contain more transcripts) than those returned by other methods, and thus fail to qualify under our notion of agreement. This occurs, partially, because `RapMap` and `Kallisto` (and to some extent `STAR`) do not tend to return sub-optimal multi-mapping locations. However, unlike `Bowtie 1`, which provided an option to return only the best “stratum” of alignments, there is no way to require that `Bowtie 2` return only the best multi-mapping locations for a read. We observe similar behavior for `Bowtie 2` (i.e. that it returns a larger set of mapping locations) in the synthetic tests as well, where the average numberofhits per read is higher than for the other methods (see Table 1). In terms of runtime, `RapMap`, `STAR` and `Bowtie 2` take 3, 26, and 1020 minutes respectively to align the reads from this experiment using 4 threads. We also observed a similar trend in terms of the average number of hits per read here as we did in the synthetic dataset. The average number of hits per read on this data were 4.56, 4.68, 4.21, 7.97 for `RapMap`, `Kallisto`, `STAR` and `Bowtie 2` respectively.

## 4 Application of quasi-mapping for transcript quantification

While mapping cannot act as a stand-in for full alignments in all contexts, one problem where similar approaches have already proven very useful is transcript abundance estimation. Recent work (Patro *et al.*, 2014; Zhang and Wang, 2014; Bray *et al.*, 2015; Patro *et al.*, 2015) has demonstrated that full alignments are not necessary to obtain accurate quantification results. Rather, simply knowing the transcripts and positions where reads may have reasonably originated is sufficient to produce accurate estimates of transcript abundance. Thus, we have chosen to apply quasi-mapping to transcript-level quantification as an example application, and have implemented our modifications as an update to the Sailfish (Patro *et al.*, 2014) software, which we refer to as quasi-Sailfish. These changes are present in the Sailfish software from version `0.7` forward. Here, we compare this updated method to the transcript-level quantification tools RSEM (Li *et al.*, 2010), Tigar2 (Nariai *et al.*, 2014) and `Kallisto` (Bray *et al.*, 2015), the last of which is based on the pseudo-alignment concept mentioned above.

### 4.1 Transcript quantification

In an RNA-seq experiment, the underlying transcriptome consists of *M* transcripts and their respective counts. The transcriptome can be represented as a set 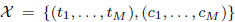, where *t_i_* denotes the nucleotide sequence of transcript *i* and *c_i_* denotes the number of copies of *t_i_* in the sample. The length of transcript *t_i_* is denoted by *l_i_*. Under ideal, uniform, sampling conditions (i.e. without considering various types of experimental bias), the probability of drawing a fragment from a transcript *t_i_* is proportional to its nucleotide fraction (Li *et al.*, 2010) denoted by 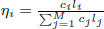.

If we normalize the η*_i_* for each transcript by its length *l_i_*, we obtain a measure of the relative abundance of each transcript called the transcript fraction (Li *et al.*, 2010), which is given by 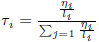.

When performing transcript-level quantification, η and τ are generally the quantities we are interested in inferring. Since they are directly related, knowing one allows us to directly compute the other. Below, we describe our approach to approximating the estimated number of reads originating from each transcript, from which we estimate τ and, subsequently transcripts per million (TPM).

### 4.2 Quasi-mapping-based Sailfish

Using the quasi-mapping procedure provided by `RapMap` as a library, we have updated the Sailfish (Patro *et al.*, 2014) software to make use of quasi-mapping, as opposed to individual k-mer counting, for transcript-level quantification. In the updated version of Sailfish, the `index` command builds the quasi-index over the reference transcriptome as described in Section 2. Given the index and a set of sequenced reads, the `quant` command quasi-maps the reads and uses the resulting mapping information to estimate transcript abundances.

To reduce the memory usage and computational requirements of the inference procedure, quasi-Sailfish reduces the mapping information to a set of equivalence classes over sequenced fragments. These equivalence classes are similar to those used in Nicolae *et al*. (2011), except that the position of each fragment within a transcript is not considered when defining the equivalence relation. Specifically, any fragments that map to exactly the same set of transcripts are placed into the same equivalence class. Following the notation of Patro *et al*. (2015), the equivalence classes are denoted as 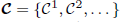, and the count of fragments associated with equivalence class C*^j^* is given by *d^j^*. Associated with each equivalence class *C^j^* is an ordered collection of transcript identifiers 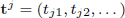 which is simply the collection of transcripts to which all equivalent fragments in this class map. We call t*^j^* the *label* of class C*^j^*.

#### 4.2.1 Inferring transcript abundances

The equivalence classes **C** and their associated counts and labels are used to estimate the number of fragments originating from each transcript. The estimated count vector is denoted by α, and α*_i_* is the estimated number of reads originating from transcript *t_i_*. In quasi-Sailfish, we use the variational Bayesian expectation maximization (VBEM) algorithm to infer the parameters (the estimated number of reads originating from each transcript) that maximize a variational objective. Specifically, we maximize a simplified version of the variational objective of Nariai *et al*. (2013).

The VBEM update rule can be written as a simple iterative update in terms of the equivalence classes, their counts, and the prior (α_0_). The iterative update rule for the VBEM is:

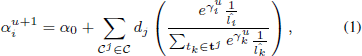

where

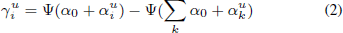

and Ψ(·) is the digamma function. Here, î*_i_* is the *effective* length of transcript *t_i_*, computed as in Li *et al*. (2010). To determine the final estimated counts — α — Equation (1) is iterated until convergence. The estimated counts are considered to have converged when no transcript has estimated counts differing by more than one percent between successive iterations.

Given α, we compute the TPM for transcript *i* as

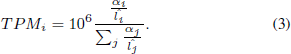

Sailfish outputs, for each transcript, its name, length, effective length, TPM and the estimated number of reads originating from it.

### 4.3 Quantification performance comparison

We compared the accuracy of quasi-Sailfish (Sailfish v0.9.0; q-Sailfish in Table 2) to the transcript-level quantification tools RSEM (Li *et al.*, 2010) (v1.2.22), Tigar 2 (Nariai *et al.*, 2014) (v2.1), and `Kallisto` (Bray *et al.*, 2015) (v0.42.4) using 6 different accuracy metrics and data from two different simulation pipelines. One of the simulated datasets was generated with the Flux Simulator (Griebel *et al.*, 2012), and is the same dataset used in Section 3 to assess mapping accuracy and performance on synthetic data. The other dataset was generated using the RSEM simulator via the same methodology adopted by Bray *et al*. (2015). That is, RSEM was run on sample `NA12716_7` of the Geuvadis RNA-seq data (Lappalainen *et al.*, 2013) to learn model parameters and estimate true expression. The learned model was then used to generate the simulated dataset, which consists of 30 million 75 bp paired-end reads.

**Table 2.** Performance evaluation of different tools along with quasi enabled sailfish (q-Sailfish) with other tools on synthetic data generated by Flux simulator and RSEM simulator.

|  | Flux simulation |  |  |  | RSEM-sim simulation |  |  |  |
| --- | --- | --- | --- | --- | --- | --- | --- | --- |
|  | Kallisto | RSEM | q-Sailfish | Tigar 2 | Kallisto | RSEM | q-Sailfish | Tigar 2 |
| Proportionality corr. | 0.74 | 0.78 | 0.75 | 0.77 | 0.91 | 0.93 | 0.91 | 0.93 |
| Spearman corr. | 0.69 | 0.73 | 0.70 | 0.72 | 0.91 | 0.93 | 0.91 | 0.93 |
| TPEF | 0.77 | 0.96 | 0.60 | 0.59 | 0.53 | 0.49 | 0.53 | 0.50 |
| TPME | -0.24 | -0.37 | -0.10 | -0.09 | 0.00 | -0.01 | 0.00 | 0.00 |
| MARD | 0.36 | 0.29 | 0.31 | 0.26 | 0.29 | 0.25 | 0.29 | 0.23 |
| wMARD | 4.68 | 5.23 | 4.45 | 4.35 | 1.00 | 0.88 | 1.01 | 0.94 |

We measure the accuracy of each method based on the estimated versus true number of reads originating from each transcript, and consider 6 different metrics of accuracy; proportionality correlation (Lovell *et al.*, 2015), Spearman correlation, the true positive error fraction (TPEF), the true positive median error (TPME), the mean absolute relative difference (MARD) and the weighted mean absolute relative difference (wMARD). Detailed definitions for the last four metrics are provided in Appendix 1.5.

Each of these metrics captures a different notion of accuracy, and all are reported to provide a more comprehensive perspective on quantifier accuracy. The first two metrics — proportionality and Spearman correlation — provide a global notion of how well the estimated and true counts agree, but are fairly coarse measures. The TPEF assesses the fraction of transcripts where the estimate is different from the true count by more than some nominal fraction (here 10%). Unlike TPEF, the TPME metric takes into account the direction of the mis-estimate (i.e. is it an over or under-estimate of the true value?). However, both metrics are assessed only on truly-expressed transcripts, and so provide no insight into the tendency of a quantifier to produce false positives.

The absolute relative difference (ARD) metric has the benefit of being defined on all transcripts as opposed to only those which are truly expressed and ranges from 0 (lowest) to 2 (highest). Since the values of this metric are tightly bounded, the aggregate metric, MARD, is not dominated by high expression transcripts. Unfortunately, it therefore has limited ability to capture the magnitude of mis-estimation. The wMARD metric attempts to account for the magnitude of mis-estimation, while still trying to ensure that the measure is not completely dominated by high expression transcripts. This is done by scaling each ARD_*i*_ value by the logarithm of the expression.

Table 2 shows the performance of all 4 quantifiers, under all 6 metrics, on both datasets. While all methods seem to perform reasonably well, some patterns emerge. RSEM seems to perform very well in terms of the correlation metrics, but less well in terms of the TPEF, TPME, and wMARD metrics (specifically in the Flux Simulator-generated dataset). This is likely a result of the lower mapping rate obtained on this data by RSEMߣs very strict `Bowtie 2` parameters. Tigar 2 generally performs very well under a broad range of metrics, and produces highly-accurate results. However, it is *by far* the slowest method considered here, and requires over a day to complete on the Flux simulator data and almost 7 hours to complete on the RSEM-sim data given 16 threads (and not including the time required for `Bowtie 2` alignment of the reads). Finally, both quasi-Sailfish and `Kallisto` perform well in general under multiple different metrics, with quasi-Sailfish tending to produce somewhat more accurate estimates. Both of these methods also completed in a matter of minutes on both datasets.

One additional pattern that emerges is that the RSEM-sim data appears to present a much simpler inference problem compared to the Flux Simulator data. One reason for this may be that the RSEM-sim data is very “clean” — yielding concordant mapping rates well over 99%, even under RSEM’s strict `Bowtie 2` mapping parameters. As such, all methods tend to perform well on this data, and there is comparatively little deviation between the methods under most metrics.

For completeness, we also provide (in Appendix 1.4) the results, under all of these metrics, where the true and predicted abundances are considered in terms of TPM rather than number of reads. We find that the results are generally similar, with the exception that TIGAR 2 performs considerably worse under the TPM measure.

## 5 Application of quasi-mapping for clustering *de novo* assemblies

Estimating gene-expression from RNA-seq reads is an especially challenging task when no reference genome is present. Typically, this problem is solved by performing *de novo* assembly of the RNA-seq reads, and subsequently mapping these reads to the resulting contigs to estimate expression. Due to sequencing errors and artifacts, and genetic variation and repeats, *de novo* assemblers often fragment individual isoforms into separate assembled contigs. Davidson and Oshlack (2014) argue that better differential expression results can be obtained in *de novo* assemblies if contigs are first clustered into groups. They present a tool, CORSET, to perform this clustering, and compare their approach to existing tools such as CD-HIT (Fu *et al.*, 2012). CD-HIT compares the sequences (contigs) directly, and clusters them by sequence similarity. CORSET, alternatively, aligns reads to contigs (allowing multimapping) and defines a distance between each pair of contigs based on the number of multimapping reads shared between them, and the changes in estimated expression inferred for these contigs under different conditions. Hierarchical agglomerative clustering is then performed on these distances to obtain a clustering of contigs.

Here, we show how `RapMap` can be used for the same task, by taking an approach similar to that of CORSET. First, we map the RNA-seq reads to the target contigs and simultaneously construct equivalence classes over the mapped fragments as in Section 4. We construct a weighted, undirected graph from these equivalence classes as follows. Given a set of contigs c and the equivalence classes *C*, we construct *G* = (*V, E*) such that V = *c*, and 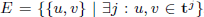. We define the weight of edge 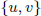 as 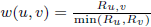. Here *R_u_* is the total number of reads belonging to all equivalences classes in which contig *u* appears in the label. *R_v_* is defined analogously. *R_u,v_* is the total sum of reads in all equivalence classes for which contigs *u* and *v* appear in the label. Given the undirected graph *G*, we use the *Markov Cluster Algorithm*, as implemented in MCL (Van Dongen, 2000), to cluster the graph.

To benchmark the time and accuracy of our clustering scheme compared to CD-HIT and CORSET, we used two datasets from the CORSET paper (Davidson and Oshlack, 2014). The first dataset consists of 231 million human reads in total, across two conditions, each with three replicates (as originally described by Trapnell *et al*. (2013)). The second dataset, from yeast, was originally published by Nookaew *et al*. (2012) and consists 36 million reads, grown in two different conditions with three replicates each. For both of these datasets, we consider clustering the contigs of the corresponding de novo assemblies, which were generated using Trinity (Grabherr *et al.*, 2011).

To measure accuracy, we consider the precision and recall induced by a clustering with respect to the true genes from which each contig originates. Assembled contigs were mapped to annotated transcripts using BLAT (Kent, 2002), and labeled with their gene of origin. A pair of contigs from the same cluster is regarded as true positive (tp) if they are from the same gene in the ground truth set. Similarly, a pair is a false positive (fp) if they are not from same gene but are clustered together. A pair is a false negative (fn) if they are from same gene but not predicted to be in the same cluster and all the remaining pairs are true negatives (tn). With these definitions of tp, fp, tn and fn we can define precision and recall in standard manner. As shown in Table 3, when considering both precision and recall, `RapMap` (quasi-mapping) enabled clustering performs substantially better than CD-HIT and similar to CORSET. `RapMap` enabled clustering takes 8 minutes and 2 minutes to cluster the human and yeast datasets respectively — which is substantially faster than the other tools. To generate the timing results above, CD-HIT was run with 25 threads. The time recorded for CORSET consists of both the time required to align the reads using `Bowtie 2` (using 25 threads) and the time required to perform the actual clustering, which is single threaded. The time recorded for `RapMap` enabled clustering consists of the time required to quasi-map the reads, build the equivalence classes and construct the graph (using 25 threads), plus the time required to cluster the graph with MCL (using a single thread). Overall, on these datasets, `RapMap`-enabled clustering appears to provide comparable or better clusterings than existing methods, and produces these clusterings much more quickly.

**Table 3.**
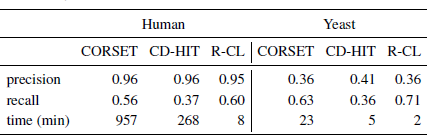
Performance of CORSET, CD-HIT and RapMap enabled clustering (R-CL) on yeast and human data.

## 6 Discussion & Conclusion

In this paper we have argued for the usefulness of our novel approach, quasi-mapping, for mapping RNA-seq reads. More generally, we suspect that read *mapping*, wherein sequencing reads are assigned to reference locations, but base-to-base alignments are not computed, is a broadly useful tool. The speed of traditional aligners like `Bowtie 2` and `STAR` is limited by the fact that they must produce optimal alignments for each location to which a read is reported to align.

In addition to showing the speed and accuracy of quasi-mapping directly, we apply it to two problems in transcriptome analysis. First, we have updated the Sailfish software to make use of the quasi-mapping information produced by `RapMap`, rather than direct k-mer counts, for purposes of transcript-level abundance estimation. This update improves both the speed and accuracy of Sailfish, and also reduces the complexity of its codebase. We demonstrate, on synthetic data generated via two different simulators, that the resulting quantification estimates have accuracy comparable to state-of-the-art tools. We also demonstrate the application of `RapMap` to the problem of clustering *de novo* assembled contigs, a task that has been shown to improve expression quantification and downstream differential expression analysis (Davidson and Oshlack, 2014). `RapMap` can produce clusterings of comparable or superior accuracy to those of existing tools, and can do so much more quickly.

However, `RapMap` is a stand-alone mapping program, and need not be used only for the applications we describe here. We expect that quasi-mapping will prove a useful and rapid alternative to alignment for tasks ranging from filtering large read sets (e.g. to check for contaminants or the presence or absence specific targets) to more mundane tasks like quality control and, perhaps, even to related tasks like metagenomic and metatranscriptomic classification and abundance estimation.

We hope that the quasi-mapping concept, and the availability of `RapMap` and the efficient and accurate mapping algorithms it exposes, will encourage the community to explore replacing alignment with mapping in the numerous scenarios where traditional alignment information is un-necessary for downstream analysis.

## Acknowledgments

The authors would like to thank Geet Duggal, Richard Smith-Unna, and Owen Dando for useful discussions regarding various aspects of this work.

## Appendix

### 1.1 Parameters for mapping and alignment tools

When `Bowtie 2` was run to produce alignment results, it was run with default parameters with the exception of `-k 200` and `-no-discordant`. When timing `Bowtie 2` the the number of threads (`-p`) was set in accordance with what is mentioned in the relevant text, and the output was piped to `/dev/null`. When `Bowtie 2` was used to produce alignment results for quantification with `RSEM, RSEM`’s `Bowtie 2` wrapper (with its default parameters) was used to generate alignemnts.

When producing alignmentresults, `STAR` was run with the following parameters: `-outFilterMultimapNmax 200 -outFilterMismatchNmax 99999 -outFilterMismatchNoverLmax 0.2 -alignlntronMin 1000 -alignlntronMax 0 -limitOutSAMoneReadBytes 1000000 -outSAMmode SAMUnosrted`. Additionally, when timing `STAR`, it was run with the number of threads (`-runThreadN`) specified in the relevant text and with the `-outSAMMode` None flag.

To obtain the “pseudo-alignments” produces by `Kallisto`, it was run with the `-pseudobam` flag.

When producing mapping results, `RapMap` was run with the option `-m 200` to limit multi-mapping reads to 200 locations. Additionally, when timing `RapMap`, it was run with the number of threads (`-t`) specified in the relevant text and with the `-n` flag to suppress output.

### 1.2 Flux Simulator parameters

The Flux simulator dataset was generated using the following parameters:

~~~
REF_FILE_NAME Human_Genome GEN_DIR protein_coding.gtf
~~~

~~~
NB_MOLECULES 5000000 TSS_MEAN 50
POLYA_SCALE NaN POLYA_SHAPE NaN
~~~

~~~
FRAG_SUBSTRATE RNA FRAG_METHOD UR FRAG_UR_ETA 350
~~~

~~~
RTRANSCRIPTION YES RT_MOTIF default
~~~

~~~
GC_MEAN NaN
PCR_PROBABILITY 0.05
PCR_DISTRIBUTION default
~~~

~~~
FILTERING YES
~~~

~~~
READ_NUMBER 150000000 READ_LENGTH 76 PAIRED_END YES ERR_FILE 76 FASTA YES
~~~

The following parameters were used to produce noise reads:

~~~
PAIRED_END YES
REF_FILE_NAME noisy.gtf
~~~

~~~
READ_LENGTH 76
PRO_FILE_NAME flux_simulator_noise_expression.pro
ERR_FILE 76
GEN_DIR Human_Genome/
SEQ_FILE_NAME noise_reads.bed
PCR_DISTRIBUTION none POLYA_SCALE NaN FASTA YES
~~~

~~~
NB_MOLECULES 2000000
READ_NUMBER 34382441
UNIQUE_IDS YES POLYA_SHAPE NaN
~~~

### 1.3 Mapping accuracy in the presence of noisy reads

We tested the effect of including background (i.e. noise) reads on the accuracy of the different mapping and alignment tools. In this experiment, we sampled 9 million reads from the 48 million read simulated data set used in Section 3.1. We then incorporated an additional 1 million “noise” reads from a simulated dataset generated with the Flux Simulator using a custom annotation. This noise annotation was created by constructing a single interval for each transcript, which contained the entire genomic range from the initial until the terminal exons (i.e. it contained all intervening intronic regions). Thus, for each annotated transcript, the noise annotation contains a nascent, un-spliced version of this transcript. This model of noise was motivated from the observation of (Gilbert *et al.*, 2004), that some RNA-seq data (e.g. human brain tissue) contains reads potentially derived from nascent, un-spliced variants of expressed transcripts.

As shown in Figure 4 we observe that, in the presence of noise, the precision for all the tools decreases slightly compared to the “clean”, 48 million read dataset described in Section 3.1. This is because some small fraction of noisy reads are assigned as false positives, as they map to the mature version of their corresponding transcript of origin that appears in the reference. Overall, however, the results follow a very similar trend both with and without noisy reads. Specifically, `RapMap` (quasi-mapping) performs almost identically to `Bowtie 2`, while `Kallisto` and `STAR` yield very similar results — somewhat under-performing `RapMap` and `Bowtie 2`. This clearly demonstrates that, in the presence of noisy reads, all of the tools degrade gracefully and still perform reasonably well, with no discernible difference between mapping and alignment-based tools.

**Fig 4:**
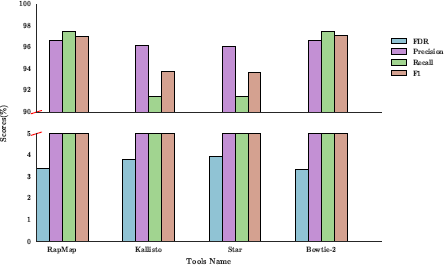
Precision, recall and F1-score (top) and FDR (bottom) on the simulated dataset with noise, for the 4 different tools we consider.

### 1.4 Quantification results using TPM

In addition to computing the error metrics based on the estimated versus true number of reads originating from each transcript (as provided in Table 2), we also evaluated the same metrics based instead on the TPM of each transcript. That is, all of the metrics defined in Section 4.3 and appendix 1.5 remain the same, except that *x_i_* now denotes the true TPM value for transcript *i* and *y_i_* denotes the estimated TPM of transcript *i*. We note that the Flux Simulator provides neither effective lengths nor TPM estimates directly. To obtain the ground truth TPM values for the Flux Simulator dataset, we first computed the effective length of each transcript (by convolving the characteristic function over the transcript with the true fragment length distribution), and then computed the TPM value for each transcript using Equation (3). The results are generally similar to what was observed at the read level, except that TIGAR 2 seems to perform considerably worse under a number of metrics on the RSEM-sim dataset when considering the TPM measure of abundance.

### 1.5 Error Metrics

We define the error metrics reported in Section 4.3 below, letting *x_i_* denote the true number of reads originating from transcript *i* and *y_i_* denote the estimated number of reads.

The relative error for transcript 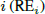 is given by 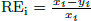 and the error indicator for transcript 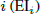 is given by

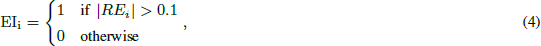

and it is equal to 1 if the estimated count for this truly expressed transcript (it is undefined, as is RE*_i_*, when *x_i_* = 0) differs from the true count by more than 10%. Given RE*_i_* and EI*_i_*, the aggregate true positive error fraction (TPEF) is defined as 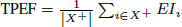. Here, *X*^+^ is the set of “truly expressed” transcripts (those having at least 1read originating from them in the ground truth). Similarly, the true positive median error is define as TPME = median 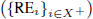.

**Table 4.** Performance evaluation of different tools along with quasi enabled sailfish (q-Sailfish) with other tools on synthetic data generated by Flux simulator and RSEM simulator.

|  | Flux simulation |  |  |  | RSEM-sim simulation |  |  |  |
| --- | --- | --- | --- | --- | --- | --- | --- | --- |
|  | Kallisto | RSEM | q-Sailfish | Tigar 2 | Kallisto | RSEM | q-Sailfish | Tigar 2 |
| Proportionality corr. | 0.79 | 0.80 | 0.80 | 0.80 | 0.94 | 0.96 | 0.94 | 0.93 |
| Spearman corr. | 0.69 | 0.73 | 0.71 | 0.60 | 0.91 | 0.93 | 0.91 | 0.89 |
| TPEF | 0.87 | 0.88 | 0.84 | 0.94 | 0.51 | 0.47 | 0.50 | 0.95 |
| TPME | 0.07 | 0.13 | 0.12 | -0.40 | 0.00 | 0.00 | 0.00 | 0.21 |
| MARD | 0.35 | 0.27 | 0.31 | 0.35 | 0.28 | 0.25 | 0.28 | 0.48 |
| wMARD | 0.67 | 1.22 | 0.69 | 1.76 | -0.74 | -0.73 | -0.74 | 0.12 |

Finally, the absolute relative difference for transcript 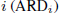 is defined as

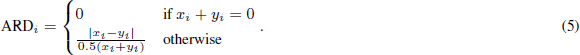

Consequently, the mean absolute relative difference (MARD) is defined as 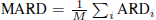, and the weighted mean absolute relative difference (wMARD) is defined as

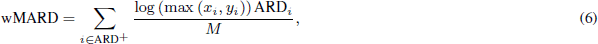

where, 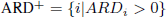, and M is the total number of transcripts.

## Footnotes

1 We do not compare against lightweight-alignment here, as no standalone implementation of this approach is currently available

